# Oncolytic Newcastle disease virus enhanced apoptosis in colorectal cancer cell lines

**DOI:** 10.1101/2023.04.16.537098

**Authors:** Teridah Ernala Ginting, Nur Rahmaniah Hidayat, Vina Cornelia, Young Othiwi Larasati, Kamaluddin Zarkasie, Irawan Yusuf

**Author notes:** corresponding author (TEG).

## Abstract

Colorectal cancer (CRC) is a deadly disease with a high prevalence and mortality rate worldwide. Previous investigations have shown that Newcastle disease virus (NDV) exhibits oncolytic activity and antitumor immunostimulation properties on several types of tumor cells but not normal cells. This study aims to examine NDV oncolytic activity against two kinds of human CRC cell lines, i.e., HCT116 and SW620, as well as its ability to induce apoptosis. The results showed that CRC cell lines were susceptible to NDV LaSota strain and the mechanism of death was due to caspase-dependent pathways apoptosis, followed by interferon signaling competence. NDV-induced proinflammatory cytokines in CRC cells might have contributed to apoptosis mechanism. Therefore, further investigation is recommended, using the findings obtained in this study as a basis for an animal CRC model.

## Introduction

Colorectal cancer (CRC) is one of the most prevalent types of cancer worldwide and often begins as polyps on the inner walls of the rectum or colon (1,2). Polypectomy during colonoscopy has been established as the primary treatment for individuals with non-metastatic colon cancer, although there are different treatment recommendations for each stage of CRC (3). Surgery or removal of part of the colon remains the standard option, despite the potential for side effects such as bleeding, infection, injury to contagious organs or failure to eliminate metastatic wounds (4). Other treatments, including chemotherapy, radiotherapy, immunotherapy, and combination therapies are also commonly used but cannot effectively combat all cancer types effectively (5). This makes it necessary to develop another therapy for CRC patients with low side effects. Oncolytic viruses have been found to selectively replicate and kill cancer without harming normal tissue. Therefore, oncolytic virotherapy is considered a promising targeted therapy against several cancer types, including CRC (6,7).

A hallmark of a viral infection is a response from infected cells, such as antiviral defense mechanism, apoptosis, and production of specific cytokines (8). Apoptosis is a vital component of various processes and a common form of cell death associated with tumor regression and immune response development (8,9). One of the methods used to detect apoptosis is the changes in phospholipid distribution in the plasma membrane. Generally, in healthy cells, phosphatidylserine is mainly located in the cytosolic side of plasma membrane and is translocated to extracellular part in early apoptosis, which can be used as a marker (10). Necrosis is an unprogrammed form of cell death that can be detected with a DNA-intercalating dye (11). Newcastle (NDV) is a negative-strand RNA virus, also known as a common disease of birds caused by avian avulavirus type 1 and a family member of **Paramyxoviridae** (12–14). NDV is a member of natural oncolytic viruses and a non-pathogenic in mammals, which can be used as a potential cancer therapy. It has oncolytic activity against mammalian cancer by immune-stimulatory capacity within human cells (15). NDV selectively infects tumor cells and the immune system responds by secreting factors that effectively kill tumors without affecting normal cells (12,16). Previous studies showed that NDV infection strongly induced the production of inflammatory cytokines such as IFN-α, IFN-β, IFN-λ, interleukin (IL)-1β, and IL-6 in many cell types (17,18). The different production of IFNs in normal and tumor cells led to the effectiveness of killing tumor cells through apoptosis (12). Other investigations reported that NDV causes apoptosis by activating the mitochondrial/intrinsic pathway and acts independently of the death receptor or extrinsic pathway (19). In this study, the potency of NDV strain LaSota against CRC cell lines Colon26, HCT116, and SW620 was investigated and the cell death, apoptosis activation, and production of the proinflammatory soluble mediator were observed. The results are expected to provide a rationale for further study in the animal CRC model.

## Materials and Methods

### Cells

Human colorectal tumor cell lines HCT116 and SW620, murine CRC Colon26, human non-malignant cell NB1RGB (skin fibroblast), HEK293 (transformed embryonic kidney), and murine transformed astroglia RCR-1 were purchased from RIKEN BioResource Center Cell Bank (Tsukuba, Ibaraki, Japan) or America Type Culture Collection (ATCC, Rockville, MD, USA). HCT116 was grown in McCoy, Colon26, and RCR-1 in RPMI-1640 (+) L-glutamine, NB1RGB cells in MEMα medium, HEK293 cells in DMEM high glucose medium, while SW620 was cultured in Leibovitz’s. All growth media were supplemented with 10% fetal bovine serum (FBS) and 1:1000 penicillin-streptomycin. Cells were cultured at 37 °C in 5% CO_2_, except for SW620 which was cultured without CO_2_. Human tumor (A549) cells (RIKEN) grown in F12-K medium containing 10% FBS were used for virus titration. Media, FBS, and antibiotics were obtained from Gibco (Carlsbad, CA, USA).

### Virus

Lentogenic NDV LaSota strain was obtained as a gift from Dr. Kamaluddin Zarkasie, from PT. IPB-Shigeta Animal Pharmaceuticals, Bogor, Indonesia. The virus was propagated in the allantoic seronegative 10-day-old embryonated chicken eggs. Allantoic fluids were collected 3 days after injection, followed by allantoic fluids centrifugation at 3000 x *g* at 4°C for 20 min and filtration with 0.45 µm to remove debris and microbial contaminant, respectively.

### Virus titration

A 96-well plate of A549 was prepared on the previous day to be confluent at the time of assay. Wells were washed with PBS (1x) pH 7.4 (Gibco) two times and a 20 μL of ten-fold serially diluted virus were added and incubated for 60 min at 37 °C in 5% CO_2_ for HCT116, or without CO_2_ for SW620. Subsequently, cells were washed twice with PBS and set in a maintenance medium that contained 4% BSA Fraction V (Gibco) and 0.1% L-(tosylamindo-2-phenyl) ethyl chloromethyl ketone (TPCK)-treated trypsin (Affymetrix, Cleveland, OH, USA). Lastly, the cells were incubated for 3 days, virus was inactivated with 10% formalin, and stained with crystal violet, and TCID_50_ calculation was performed according to Reed and Muench (20).

### NDV cytotoxicity in CRC cell lines

MTT assay was used to examine the cytotoxicity of CRC cells after NDV infection. Tetrazolium salt or 3-(4,5-dimethylthiazole-2-yl)-2,5-diphenyltetrazolium bromide) was used to measure cell proliferation and cytotoxicity. HCT116 and SW620 were plated separately overnight in a 96-well plate at a density of 5 × 10^4^ cells. Similarly, normal cells HEK293, NB1RGB, and RCR-1 were also prepared, followed by infection of confluent cells with NDV 0.001 MOI, which were then incubated for 24, 48, and 72 hours. After incubation, 20 µL of 5 mg/mL MTT substrate was added and plates were incubated for 4 hours at 37°C. Subsequently, the purple crystal formed was dissolved with Dimethylsulfoxide (DMSO) (Sigma), and the absorbance was read at 540 nm. Data were presented as a percentage of cells viability.

### NDV infection to investigate the cytokine production

To determine the cytokine production of cells after NDV infection, confluent 8 × 10^5^ HCT116 and SW620 cells per well were prepared in 6-well plates. Cells were washed twice with PBS and plates were incubated with 200 μL virus in maintenance medium at 0.001 and 0.0001 MOI, at 37°C in 5% CO_2_ for 60 min for HCT116 cells, while SW620 cells were incubated in the same condition but without CO_2_. Negative control was prepared by incubating cells with 200 μL PBS only. After incubation, cells were washed three times with PBS to remove unbound virus and a 2 mL maintenance medium was added to each well. Culture supernatants were collected at 24, 48, and 72 hours after infection for further assay.

### Multiplex proinflammatory cytokines: Enzyme-Linked Immunosorbent Assay (ELISA)

To determine proinflammatory cytokines in culture supernatant post-NDV infection, Veriplex® 9-plex human cytokine ELISA kit (PBL Assay Science, Piscataway, NJ, USA) was used according to the manufacturer’s protocol. Cell supernatant after 48 hours of NDV 0.001 and 0.0001 MOI infections was used in this assay. Collected supernatants were UV-inactivated for 30 min before assay and diluted in a 1:1 buffer solution. This was followed by incubation with coated antibodies against 9 different proinflammatory cytokines (type I interferon (IFN-α, IFN-β, and IFN-ω), type II interferon (IFN-γ), type III interferon (IFN-λ1/2/3), and other proinflammatory cytokines, including TNF-α, IP-10, IL-1-α, and IL-6, for 2 hours at room temperature. Plate was washed and 50 µL of biotinylated detection antibody was added into each well for 1 hour at room temperature. Subsequently, a 50 µL chemiluminescent substrate was added to each well and incubated for 15 minutes at room temperature. Chemiluminescence intensity was visualized using VersaDoc 4000 imaging system (BioRad, Hercules, CA, USA) and analyzed with Q-View® software (Quansys Bioscience, Logan, UT, USA). Data were collected from triplicate wells and graphs presented average pixel intensity.

### Annexin-V and PI staining flow cytometry

Flow cytometry was carried out using propidium iodide (PI) with annexin-V to investigate the cytotoxicity of NDV and cell death mechanism in CRC cells HCT116, SW620, and Colon26. Confluent cells were infected with NDV 0.001 and 0.01 MOI and prepared as a single-cell suspension using Accutase (Gibco). Cells were stained with antibodies annexin-V–FITC (Cell Signaling Tech, Massachusetts, USA), followed by PI–alexa488 (Cell Signaling). Then cells were fixed with 2% formalin for 20 min and analyzed on a FACS Accuri C6 flow cytometer (Becton-Dickinson, Franklin Lake, NJ, USA). Flow cytometry analyses were performed with BD CSampler(tm) Software (Becton-Dickinson, Franklin Lake, NJ, USA) and graphs were obtained from average fluorescence intensity.

### Caspase-3 enzyme activity assay in NDV-infected cells

Confluent cells were infected with NDV 0.001 and 0.0001 MOI for 24 hours. A positive control was prepared by treating cells with 1.0 μM staurosporine, which induces apoptosis and a negative control was prepared as mock-infected cells. The next day, cells were extracted in RIPA buffer (Sigma) supplemented with protease and phosphatase inhibitors. Proteins were collected by centrifugation at 12,000 rpm for 20 min at 4°C, run on 12% acrylamide gels at 110 V for 60 min, and transferred into 0.2 μM nitrocellulose membrane in transfer buffer containing tris 25 mM, glycine 192 mM, methanol 20%, and H2O, at 25 V for 30 min. Furthermore, HCT116 lysate was prepared to be visualized by chemiluminescence imaging. In this study, we could not get enough SW620 lysate and the western blot signal was too weak to detect by chemiluminescence, therefore, SW620 lysate was subjected for fluorescence imaging. Briefly, membranes were washed with TBS and incubated in a blocking buffer for 60 min at room temperature. For HCT116, the blocking buffer was 0.5% skim milk in TBS, while for SW620, it was Odyssey blocking buffer TBS (Li-Cor 927-60001) for 60 min. Membranes were incubated overnight at 4°C with 1:1000 diluted primary anti-caspase-3, human/mouse caspase-3 antibody polyclonal goat IgG (R&D systems) for HCT116, or caspase-3 antibody monoclonal mouse IgG (Abcam) for SW620. Next, HCT116 membrane was incubated in HRP-conjugated secondary antibody diluted 1:5000, while SW620 was incubated in IRDye-conjugated secondary antibody diluted 1:15000. Secondary antibodies were diluted in corresponding blocking buffer. HCT116 membrane was then incubated with ELISABright chemiluminescent substrate (Advansta, Bering Drive, Sa Jose, USA) and visualized using VersaDoc imaging system (BioRad), while SW620 membrane was visualized using Odyssey CLx (LiCor).

### Chromogenic in situ hybridization (ISH)

To confirm the infectivity of NDV, an ISH assay was performed to detect NDV mRNA transcripts in cells using a commercial kit RNAscope Brown Assay (Advanced Cell Diagnostics, Newark, USA) according to the manufacturer’s instructions. CRC cells were cultured on a sterile, poly-L-Lysine (Sigma) coated coverslip, and confluent cells were treated with NDV 0.001 MOI and 0.0001 MOI for 24 hours. The next day, cells were fixed with 10% neutral buffered formalin (NBF), followed by incubation in an increasing alcohol bath of 50%, 70%, and 100%. Cells were treated with RNAscope hydrogen peroxide for 10 min at RT, rinsed with PBS twice, followed by incubation in RNAscope Protease III for 10 min at RT. The assay began with hybridization using an NDV detection probe and amplification using probes 1 to 6 provided in the kit and continued with staining using hematoxylin and eosin staining. Images were acquired with a Zeiss Axioskop 40 microscope (Carl Zeiss Microimaging, Göttingen, Germany).

## Results

### NDV did not replicate efficiently in CRC cell lines

Virus particles were measured using TCID_50_ assay to determine the replication of NDV in CRC HCT116, SW620, and Colon26. TCID_50_ is based on the end-point dilution of virus at which a (CPE) is detected in 50% of the cell culture replicates infected by the given amount of virus suspensions (20). The measurement of infectious virus particles by a TCID_50_ assay showed that NDV at a concentration of 0.001 MOI did not replicate in CRC cell lines. Therefore, NDV with 0.001 MOI concentration was selected to be a dose standard for further experiments.

**Table 1.** Replication of NDV in colorectal carcinoma cells using TCID_50_

| Cell name | Tissue | Log<sub>10</sub> TCID<sub>50</sub> hours p.i. |  |  |
| --- | --- | --- | --- | --- |
|  |  | 24 | 48 | 72 |
| HCT116 | Colorectal carcinoma | <1 | <1 | <1 |
| SW620 | Colon adenocarcinoma | <1 | <1 | <1 |
| Colon26 | Murine colorectal carcinoma | <1 | <1 | <1 |

### NDV RNA expression detection in CRC cells using ISH RNAscope

ISH RNAscope was used to visualize NDV mRNA in HCT116, SW620, and Colon26 CRC cell lines after NDV infection. The brown punctate staining pattern represented a single NDV mRNA transcript in CRC and was detected in NDV-infected cells but not in uninfected control cells. The cells were stained with a probe against the rat peptidylprolyl isomerase B (PPIB) gene as a positive control probe, and *Bacillus subtilis* dihydrodipicolinate reductase (dapB) gene as a negative control probe to assess the assay specificity and RNA integrity. The number of NDV mRNA transcripts was counted by examining quantitative chromogenic spots in randomly selected cells in microscopic images. The intensity of NDV mRNA transcript dots was based on levels of NDV in infected cells, where more intense brown dots indicated a higher number of transcripts in the cells. The results of RNAScope showed brown dots that revealed the presence of NDV mRNA transcripts in NDV-infected CRC cells at a dose of 0.001 MOI, indicating NDV replication in the cytoplasm of infected cells. Fig 1 demonstrated that NDV-infected cells showed abundant, punctate brown dots, and the number and intensity were MOI-dependent. Furthermore, there was no NDV signal detected in uninfected cells.

**Fig. 1.**
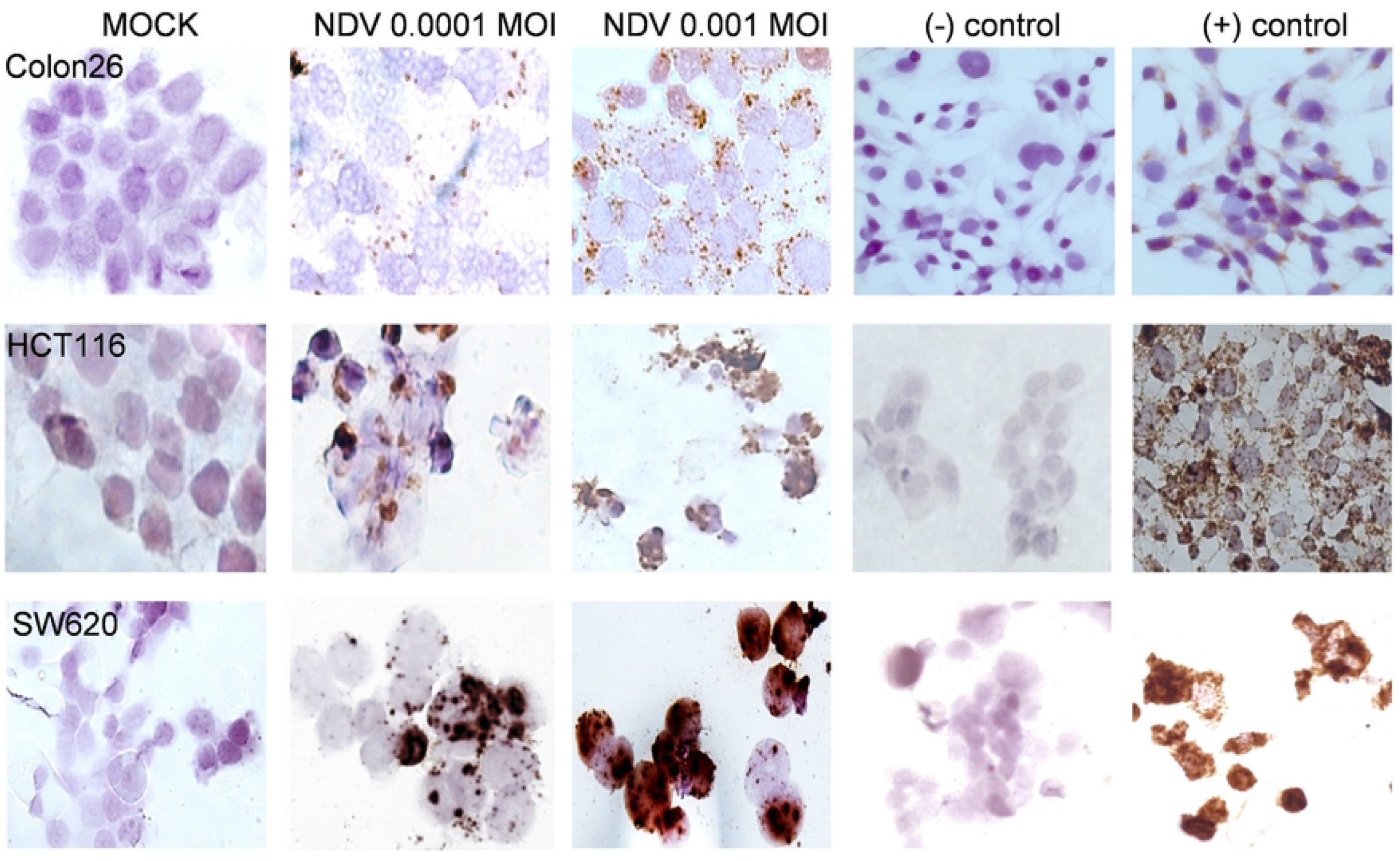
NDV mRNA transcription on CRC cells HCT116, SW620, and Colon26. RNAscope brown assay specifically detected NDV mRNA expression in CRC. NDV-infected cells showed punctate brown staining with RNAscope brown assay, while controls did not. The experiments were performed in duplicates.

### NDV infection reduced the viability of CRC cells

To determine the viability of CRC cells after NDV infection at 0.001 MOI, MTT assay was carried out on CRC cell lines HCT116, SW620, and Colon26. As a comparison in normal cells, similar experiments were conducted on normal HEK293, NB1RGB, and RCR-1 cell lines. Fig 2A showed that NDV treatment on CRC cells decreased cell viability significantly. Meanwhile, normal cells in Fig 2B were relatively resistant to NDV infections and survived at the end of observation. The results showed that colorectal cells HCT116 SW620 and Colon26 were susceptible to NDV infection compared to normal cells.

**Fig. 2.**
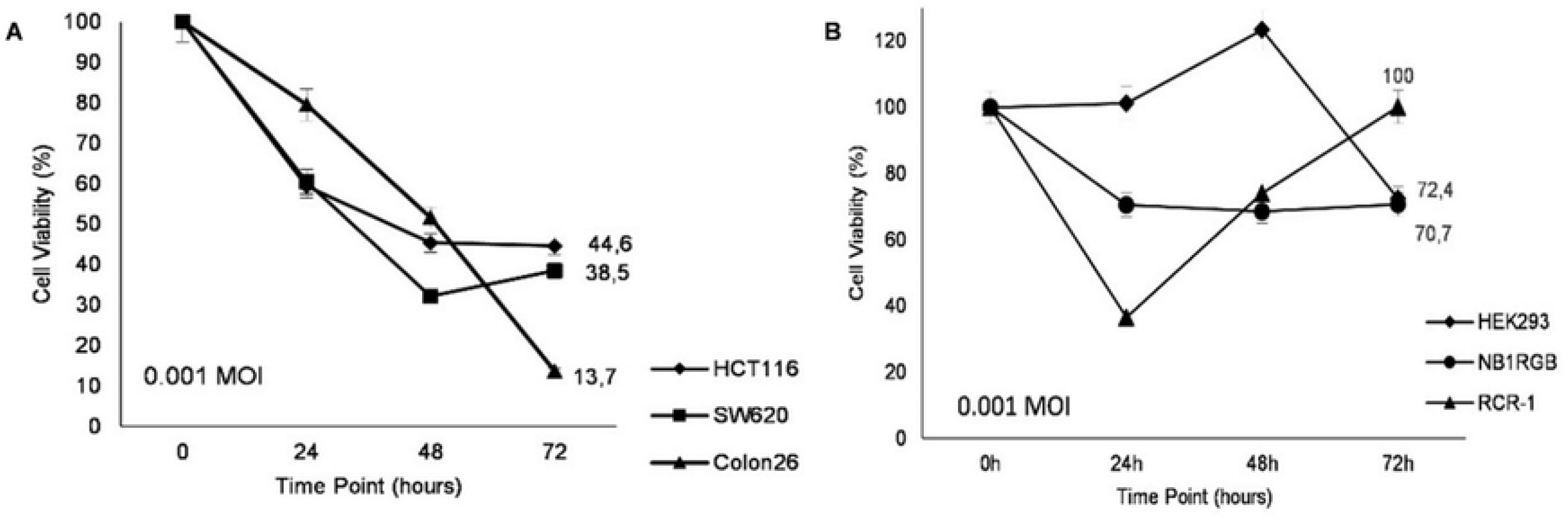
Cytotoxicity of NDV in CRC (HCT116, SW620, and Colon26) and normal cells (NB1RBG, RCR-1, and HEK-293). (A) NDV was cytotoxic to CRC cells, (B) while normal cells are relatively resistant. Percentage of viability cells normalized to mock-infected cells and presented from the ratio of treated cells with mock-infected cells. Data were collected from triplicate wells.

### NDV infection in CRC induced the apoptotic pathways

Apoptotic activity was assessed to determine whether CRC cell death induced by NDV was due to apoptosis. This was performed by measuring the levels of annexin-V and caspase-3 using flow cytometry and western blot, respectively. Annexin-V and PI were used to detect the cell membrane’s phosphatidyl serine exposure and PM integrity, respectively. Caspase are a family of proteins that are activated in the early stages of apoptosis and are an essential hallmark for apoptotic chromatin condensation and DNA fragmentation in all cell types (21).

HCT116 and SW620 were treated with increasing concentrations of NDV at doses 0.01 and 0.001 MOI and the percentage of apoptotic cells was determined with annexin-V/PI staining. The results of flow cytometry (Fig 3) revealed that both CRC cells induced apoptotic following NDV treatment, although each cell type responded differently. Compared to SW620, HCT116 was more reactive to NDV treatment because it induced more apoptosis, indicating that the effects of NDV infection on solid cancer are more effective than metastatic cancer. CRC cells were stimulated with 1 mM staurosporine for positive control staining of apoptosis, while for the negative control, cells were untreated.

**Fig. 3.**
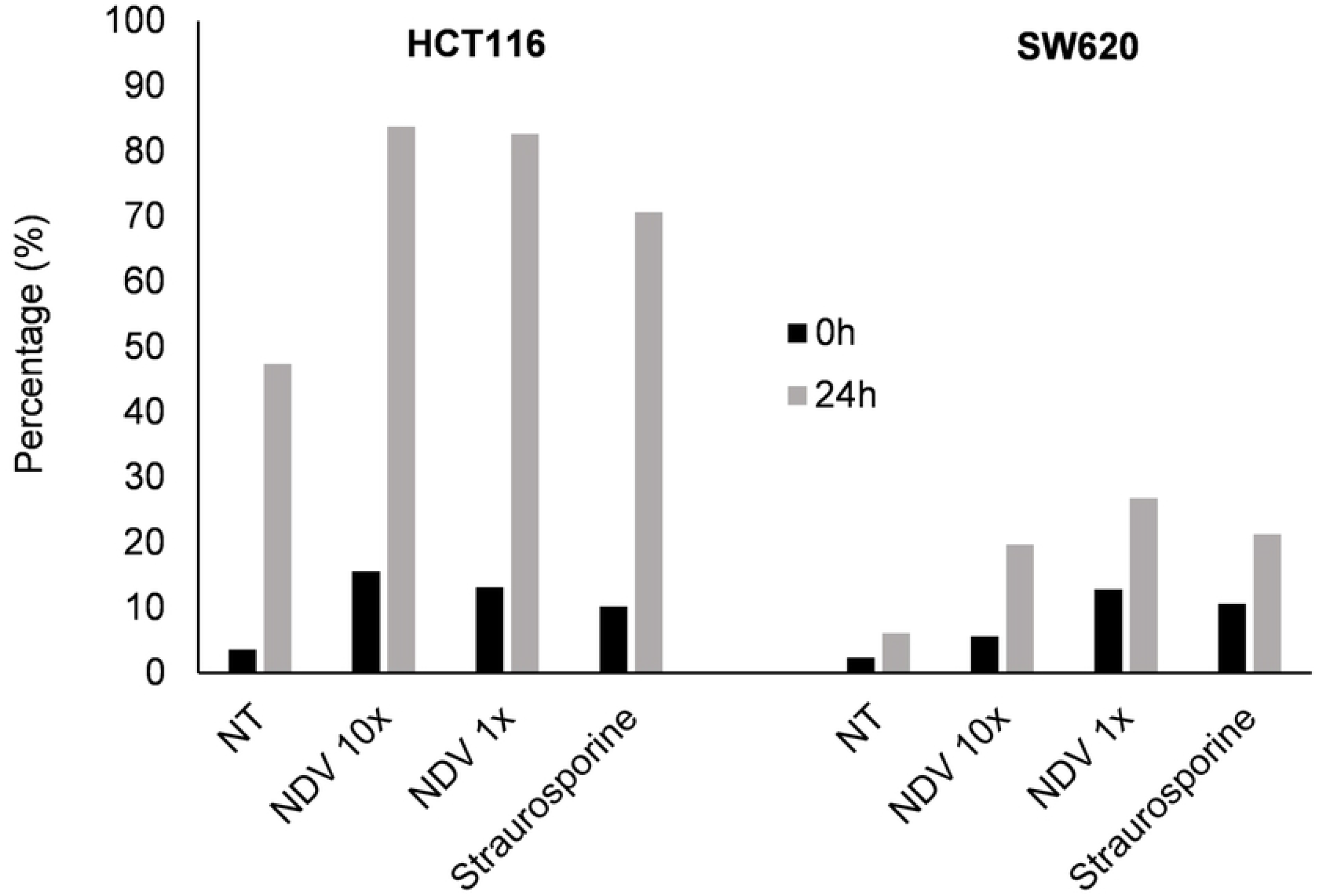
Annexin-V expression in HCT116 and SW620 after NDV infection (0h and 24h). Annexin-V/propidium iodide was performed to detect cell apoptosis. The graph showed the percentage of the apoptotic human colorectal carcinoma cells, HCT116 and SW620, at indicated MOI of NDV. The experiments were performed in duplicate.

A previous study showed that proinflammatory responses induced by NDV infection in certain tumor cells contribute to tumor-selective apoptosis. Therefore, to determine whether CRC cytotoxicity by NDV infection is due to apoptosis, HCT116, SW620, and Colon26 were infected with NDV at 0.001 and 0.0001 MOI, while caspase-3 activity and proinflammatory cytokine in cells was examined after 48 hours of infection. The cells after 48 hours were selected because they showed a distinct reduction of viability, which might be the consequence of cytokine expression. Next, we demonstrated caspase-3 activity with western blotting using specific antibodies against caspase-3 substrates (22) in cell lysate (Fig 4). The results showed NDV-induced upregulation of caspase-3 activity in HCT116 and SW620 cells. It was shown that NDV might induce cell death through the activation of caspase-3 pathway. However positive control staurosporine did not induce caspase-3 cleavage in SW620, which was consistent with a previous study in colorectal carcinoma cells SW480 (23).

**Fig. 4.**
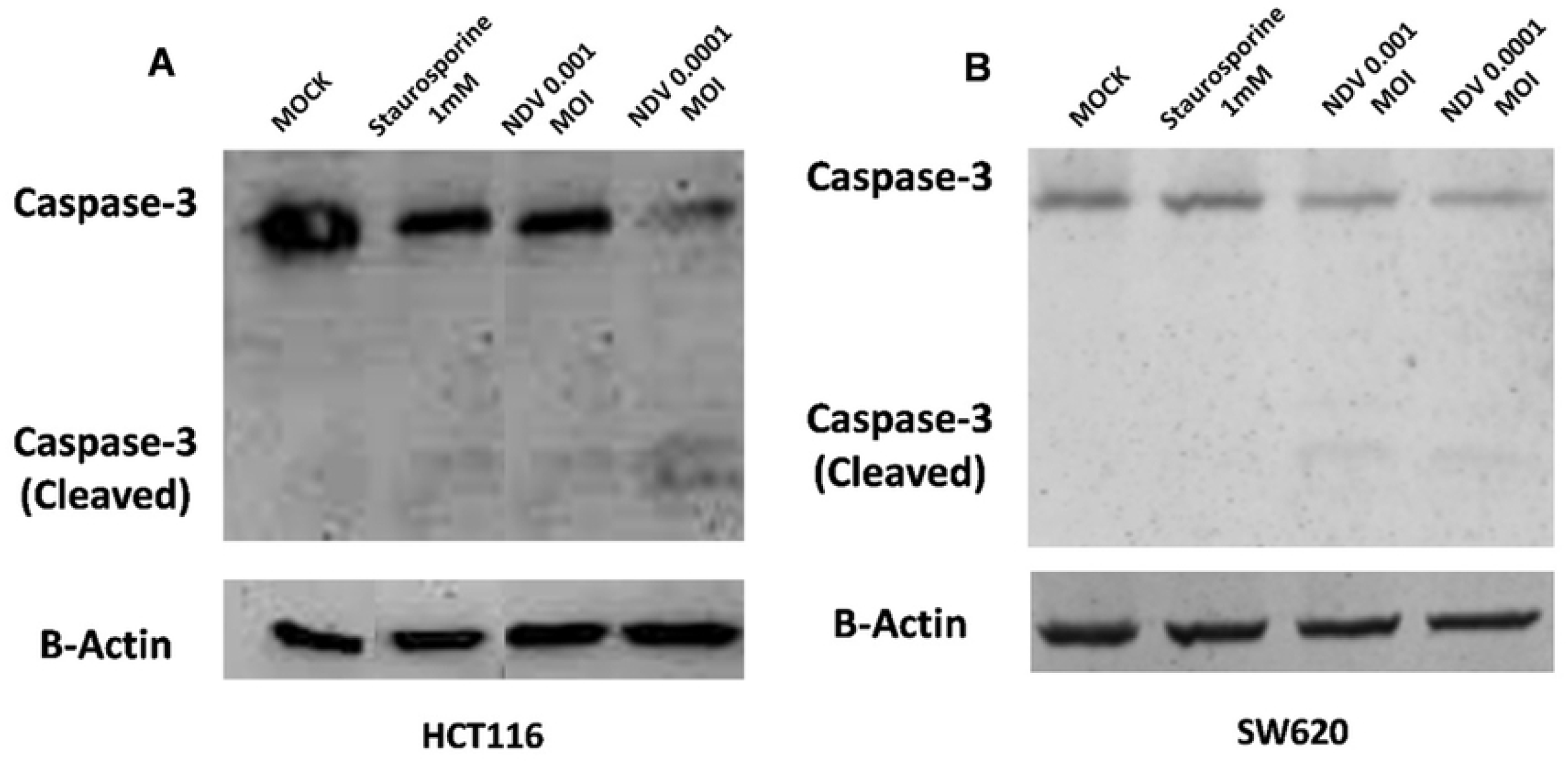
Caspase-3 activity in CRC. Caspase-3 activity in CRC cells was determined by western blot. (A-B) Western blot showed detected expression of caspase-3 and cleaved caspase-3 in HCT116 and SW620. Stausporine 1 mM was used as a positive control for apoptotic cells.

### Proinflammatory cytokines response in CRC cell after NDV infection

One of the characteristics of cancer cells is the deficiency in interferon (IFN) pathway (24), along with other tumor cells, which makes them sensitive to NDV. NDV stimulates the immune system to produce proinflammatory cytokines with antitumor capacities, such as various IFN. However, a viral infection of tumors can promote immunosuppressive surroundings by inducing immune cytokines and chemokines (25).

A multiplex analysis of proinflammatory cytokines Veriplex Human Interferon 9-Plex Elisa was conducted to determine the roles of NDV infection in inflammatory response induction in CRC cell lines. The results showed that NDV infection strongly induced proinflammatory cytokines IFN-α, IFN-β, IFN-λ, and IP-10 in CRC cell lines (Fig 5). However, each CRC, i.e. HCT116 or SW620, responded differently toward NDV infection. HCT116-infected NDV secreted primarily type I IFN (IFN-α, IFN-β), type III IFN (IFN-λ), and IP-10, while SW620 responded to NDV by secreting IFN-λ and IP-10. IFN-gamma-inducible protein (IP-10) was known as a biomarker of viral infection and had been reported to be produced by airway epithelium upon infection with respiratory viruses (17). Level of IP-10 correlated with a viral titer (16), hence, NDV-induced expression of IP-10 was dependent on interferon induction.

**Fig. 5.**
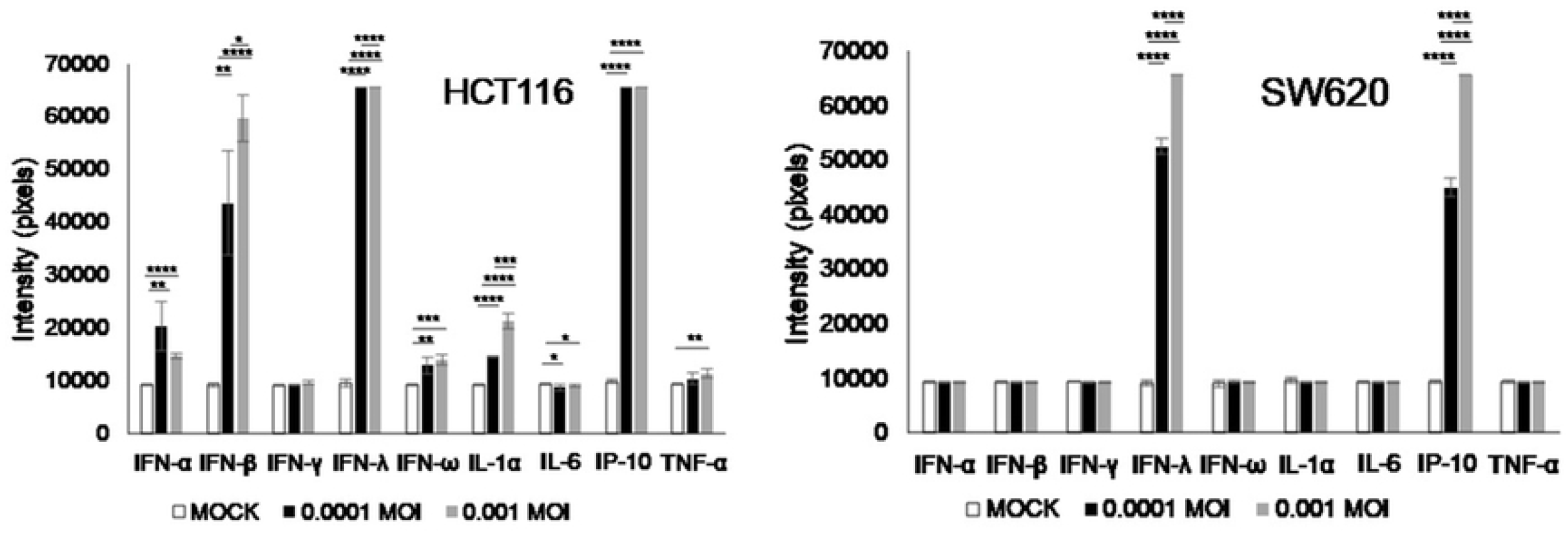
Proinflammatory cytokine-induced by NDV (0.001 and 0.0001 MOI). The graph showed the intensity of proinflammatory cytokine in the supernatant of HCT116 and SW620 induced by NDV 0.001 and 0.0001 MOI.

## Discussion

The high mortality rates associated with CRC necessitate the development of new cancer therapies with minimal consequences. In this study we provide evidence that NDV is a potent oncolytic virus in CRC cells by apoptosis. This study aimed to provide basic evidence of NDV oncolytic for CRC therapy in the laboratory for further animal and clinical study.

The results showed that NDV LaSota strain can infect CRC cell lines HCT116 and SW620, as indicated by the presence of NDV mRNA transcript in cells by ISH but the virus was not efficiently released to culture supernatant. Cytotoxicity studies showed that CRC cell lines were susceptible to NDV infection at 0.001 and 0.0001 MOI compared to normal cells. This suggested that NDV has the potential as an oncolytic agent against CRC cells while being safe for normal cells. The mechanism of CRC cell death induced by NDV was due to apoptosis based on annexin-V flow cytometry and caspase-3 activity assay. Apoptosis induction observed was more remarkable in HCT116 compared to SW620 and no cleaved caspase-3 was detected in SW620 treated by staurosporine. Similarly, Malsy et al., (2019) reported that staurosporine stimulation did not induce apoptosis in colorectal carcinoma cells SW480. However, the signaling pathway by which staurosporine induces apoptosis remains unclear. This makes it necessary to further investigate the differences in the mechanism of apoptosis between HCT116 and SW620 due to their various nature, which were solid and metastatic tumors, respectively.

This study demonstrated that NDV infection activated a robust immune response in CRC cell lines as indicated by the increase of IFN-α, IFN-β, IFN-λ, and interferon gamma-induced protein 10 (IP-10), although both CRC cells responded differently toward NDV. Interferon has been shown to have antiviral, anti-proliferative, and immunomodulatory effects in the host response to viral or bacterial infection. Based on NDV infections, previous investigations reported that IFN-α, IFN-β, and IFN-λ induced by NDV contributed to the high level of apoptosis in tumor cells.

## Conclusion

This study demonstrated that NDV LaSota can infect CRC cell lines, specifically HCT116 and SW620, without releasing its progeny virus. Compared to normal cells, CRC cell lines were more susceptible to NDV infection. The oncolysis was induced by the higher production of proinflammatory cytokines, IFN-λ, and IP-10, in both CRC cell lines. These cytokines possibly contributed to cell death through apoptosis, as indicated by the upregulation and activation of caspase-3 in both CRC cell lines. However, apoptosis was higher in HCT116 compared to SW620 cells. Further investigation is required to explore the different mechanisms of cell death in SW620 cells.

The results showed that NDV treatment can potentially suppress the growth of CRC cells through apoptosis and the production of proinflammatory cytokines. This suggested the potential of NDV as an oncolytic agent in non-metastatic CRC. Moreover, further investigations are needed to explore the potency of NDV oncolytic in CRC animal models.

## Acknowledgments

This study was supported by the Mochtar Riady Institute for Nanotechnology.

